# The role of recombination on genome-wide patterns of local ancestry exemplified by supplemented Brook Charr populations

**DOI:** 10.1101/702696

**Authors:** Maeva Leitwein, Hugo Cayuela, Anne-Laure Ferchaud, Éric Normandeau, Pierre-Alexandre Gagnaire, Louis Bernatchez

## Abstract

Assessing the immediate and long-term evolutionary consequences of human-mediated hybridization is of major concern for conservation biology. Several studies have documented how selection in interaction with recombination modulates introgression at a genome-wide scale, but few have considered the dynamics of this process within and between chromosomes. Here, we used an exploited freshwater fish, the Brook Charr (*Salvelinus fontinalis*) for which decades of stocking practices have resulted in admixture between wild populations and an introduced domestic strain to assess both the temporal dynamics and local chromosomal variation in domestic ancestry. We provide a detailed picture of the domestic ancestry patterns across the genome using about 33,000 mapped SNPs genotyped in 611 individuals from 24 supplemented populations. For each lake, we distinguished early and late-generation hybrids using admixture tracts information. To assess the selective outcomes following admixture we then evaluated the relationship between recombination and admixture proportions at three different scales: the whole genome, chromosomes and within 2Mb windows. This allowed us to detect the signature of varied evolutionary mechanisms, as reflected by the finding of genomic regions where the introgression of domestic haplotypes are favored or disfavored. Among these, the main factor modulating local ancestry was likely the presence of deleterious recessive mutations in the wild populations, which can be efficiently hidden to selection in the presence of long admixture tracts. Overall, our results emphasize the relevance of taking into consideration local ancestry information to assess both the temporal and chromosomal variation in local ancestry toward better understanding post-hybridization evolutionary outcomes.

## Introduction

Understanding the evolutionary consequences of voluntary or involuntary anthropogenic hybridization is of major concern for conservation biology (Waples, 1991; Allendorf, 2017; McFarlane & Pemberton, 2018). Indeed, anthropogenic hybridization may affect fitness components (survival, growth and reproduction), which may in turn impact population dynamics, genetic diversity and long-term viability (e.g. Allendorf, Hohenlohe, & Luikart, 2010; McFarlane & Pemberton, 2018). However, the consequences of induced gene flow between foreign and local populations are not well understood and have been considered as potentially either beneficial or harmful to the local populations, depending on the context (e.g. Todesco et al., 2016; McFarlane & Pemberton, 2018). On the positive side, the introduction of foreign individuals may be used to rescue endangered, inbred populations (i.e. genetic rescue) with the goal of increasing the mean fitness of individuals in the local population (Frankham, 2015; Harris, Zhang, & Nielsen, 2019). On the negative side, outbreeding depression may occur when the extant of genetic divergence between populations or species is sufficiently important to cause genetic incompatibilities (i.e. Dobzhansky-Muller Incompatibilities; Orr, 1995; Turelli & Orr, 2000), which may lead to a loss of local adaptation through the disruption of co-adapted genes (Waples, 1991; Verhoeven, Macel, Wolfe, & Biere, 2011). Additionally, while positive effects may be observed in the first hybrid generations by masking the effect of accumulated recessive deleterious alleles (i.e. associative overdominance) (Lippman & Zamir, 2007; Chen, 2010; Kim, Huber, & Lohmueller, 2018; Harris et al., 2019), negative effects may arise in later generations of admixture when maladapted recessive alleles are exposed to selection (Racimo, Sankararaman, Nielsen, & Huerta-Sánchez, 2015; Harris & Nielsen, 2016; Harris et al., 2019).

Therefore, considering both the time since hybridization has occurred and the recombination rate variation along the genome is critical to distinguish between the immediate and long term consequences of admixture (Harris & Nielsen, 2016; Harris et al., 2019). Since recombination is expected to progressively reduce the length of introgressed haplotypes across generations following initial admixture (Racimo et al., 2015), the length of introgressed haplotypes can be used as a proxy to estimate the time since hybridization (Gravel, 2012; Racimo et al., 2015). Thus, longer admixture tracts are expected in early hybrids while later hybrid generations tend to display shorter tracts (Racimo et al., 2015; Leitwein, Gagnaire, Desmarais, Berrebi, & Guinand, 2018). Additionally, the local variation in the recombination rate is also expected to affect the introgressed haplotype length with longer and shorter haplotypes expected in lower and higher recombining regions, respectively (S. Martin & Jiggins, 2017; Racimo et al., 2015). As a consequence, the effects of selection in interaction with recombination should vary along the genome between low and high-recombination regions. Thus, in genomic regions of low recombination, long introgressed haplotypes may totalize the individual effects of multiple selected mutations acting collectively at a block scale (*sensus* Anderson & Stebbins, 1954), as in early-generation hybrids (Leitwein et al., 2018). One could expect such block effect to generate lower introgression rate in low recombining regions due to genetic incompatibilities or barrier to introgression (Schumer et al., 2018; S. H. Martin, Davey, Salazar, & Jiggins, 2019). Such pattern was observed in swordtail fish hybrid populations for which the introgressed ancestry was more persistent in high recombining regions where incompatibility alleles uncoupled more quickly. Inversely, higher introgression rate can also be observed in low recombining regions due to the presence of recessive deleterious mutations (i.e. associative overdominance; Kim et al., 2018; Harris et al., 2019). This is because longer haplotypes will be more efficient for masking the effect of multiple recessive deleterious alleles (S. Martin & Jiggins, 2017; Racimo et al., 2015; Leitwein et al., 2018). In genomic regions of high recombination rate but also in later hybrid generations, introgressed haplotypes should be shorter and thus selective effects would be more likely to be revealed locally, that is at the locus scale. Therefore, both highly recombining regions and anciently introgressed haplotypes could display either low or high introgression rates depending on the adaptive or maladaptative nature of introgressed alleles at specific loci and the mutation load of the recipient populations (e.g. Harris & Nielsen, 2013; Racimo et al., 2015; Harris & Nielsen, 2016; Harris et al., 2019).

Clearly, variable patterns of genome-wide admixture and introgression may result from the interplay of multiple evolutionary processes (e.g., drift, positive or negative selection for the introgressed alleles) rather than a single, general mechanism. Moreover, antagonistic evolutionary mechanisms (e.g., positive and negative selection) may act differentially across the genome, especially if several pulses of hybridization have occurred, resulting in both historical and contemporary gene flow within a population from an exogenous source (Gravel, 2012; McFarlane & Pemberton, 2018). A few recent studies have investigated how the interaction between recombination rate and selection may modulate the genome-wide temporal dynamics of introgression (e.g. Martin & Jiggins, 2017; Duranton et al., 2018; Kim et al., 2018; Schumer et al., 2018; Harris et al., 2019; Martin et al., 2019). Even fewer studies have empirically investigated the temporal dynamics of introgression at the local genomic scale with the general goal of testing the above, alternative expectations (but see Martin et al., 2019 for a chromosomal approach).

The general goal of this study was to investigate the level of heterogeneity of admixture along the genome, as well as the role of mechanisms underlying those variations in a freshwater fish, the Brook Charr. In Québec Canada, this socio-economically important species has undergone intense stocking from a domestic strain for many decades (detailed in Létourneau et al. 2018). The history of each stocking event has been recorded in provincial wildlife reserves, which allows assessing the temporal dynamics of the domestic introgression in wild populations (Lamaze, Sauvage, Marie, Garant, & Bernatchez, 2012; Létourneau et al., 2018). In a recent study, Létourneau et al. (2018) documented a negative relationship between the proportion of domestic ancestry and the mean number of years since the most important stocking event (Létourneau et al., 2018). However, this study did not investigate the selective consequences (positive or negatives) of the introgressed domestic ancestry within wild populations which thus remains poorly understood. The recent availability of a high density linkage map developed for *S. fontinalis* (Sutherland et al., 2016) and a reference genome for the sister species the Arctic Charr (*Salvelinus alpinus*) (Christensen et al., 2018) open new opportunities to investigate for the first in any salmonid how selection in interaction with recombination modulates introgression at a genome-wide scale, as well as within and between chromosomes following human-mediated hybridization events.

More specifically, we used a RADseq data set collected from 24 Brook Charr populations that have been stocked with the afore mentioned domestic strain to assess genome-wide patterns of variation in local domestic ancestry for both early and late-generation hybrids. Moreover, we considered the local recombination rate to investigate which selective effects (positive, negative or neutral) may drive the genome-wide domestic ancestry pattern at three different scales; whole genome, chromosomes and 2Mb sliding windows size. We finally, examined how the presence of putative deleterious mutations may modulate the genome-wide domestic ancestry, also taking recombination rate into account.

## Materials and Methods

### Study system

As many salmonids, the Brook Charr is a socio-economically important species that is highly valued for recreational fishing. As a consequence, intensive stocking programs have been developed to support this industry. The domestic Brook Charr strain has been reproduced and maintained in captivity for more than 100 years to sustain supplementation programs (Ministère du Développement Durable, de l’Environnement, de la Faune et des Parcs 2013). On average, in the province of Québec, Canada, more than 650 tons of Brook Charr are released annually into the wild (Ministère du Développement Durable, de l’Environnement, de la Faune et des Parcs 2013; Létourneau et al., 2018) resulting in frequent hybridization between wild and domestic populations (Marie et al. 2010; Lamaze et al., 2012; Létourneau et al., 2018).

### Sampling, sequencing and genotyping

The Brook Charr populations analyzed in this study were sampled in 2014 and 2015 (Létourneau et al., 2018) and consist in 611 individuals from 24 lakes located in two wildlife reserves (Mastigouche and St-Maurice) in Québec, Canada. Additionally, 37 domestic fish originating from the *Truite de la Mauricie* Aquaculture Center broodstock, were used as reference for domestic samples. Stocking intensity was variable among lakes,as can be seen from the data on the history of stocking including the number of years since the mean year of stocking (mean_year), the total number of stocking events (nb_stock_ev), the mean number of fish stock per stocking event (mean_stock_fish) and the total number of fish stocked per stocking event (total_ha) reported in Table 1. GBS library preparation was performed in Létourneau et al. (2018), after the extraction of genomic DNA from fin clips and quality evaluation. Libraries were amplified by PCR and sequenced on the Ion Torrent Proton P1v2 chip. Raw reads were checked for quality and the presence of adapters with FastQC (http://www.bioinformatics.babraham.ac.uk/projects/fastqc/), reads were then demultiplexed with STACKS v1.40 (Catchen, Hohenlohe, Bassham, Amores, & Cresko, 2013) with the option *process_radtags* as described in Létourneau et al. (2018). Demultiplexed reads were aligned to the Arctic Charr (*Salvelinus alpinus*) reference genome (Christensen et al., 2018) with BWA_mem program v. 0.7.9 (Li & Durbin, 2010) before the individual SNPs calling with *pstacks* module (using m=3 and the bounded error model with α=0.05). To build the catalogue in *cstacks*, we randomly used 10 individuals per population with a coverage depth of at least 10X and a minimum of 500,000 reads. Each individual was then matched against the catalogue with *sstacks*. The *population* module was then run separately for each of the 24 lakes and the domestic strain, in order to generate one VCF file per population with loci passing the following filters: (i) a minimum depth of 4 reads per locus, (ii) a genotype call rate of at least 60% per population, (iii) a minimum allele frequency of 2% and (iv) a maximum observed heterozygosity of 80%. Individuals with a high percentage of missing data (>20%) were removed resulting in a final data set of 33 domestic individuals and 603 wild caught individuals (Table 1). Finally, to avoid merging paralogs, we removed for each individual loci with more than two alleles with the R package *stackr* (Gosselin & Bernatchez, 2016).

**Table 1.**
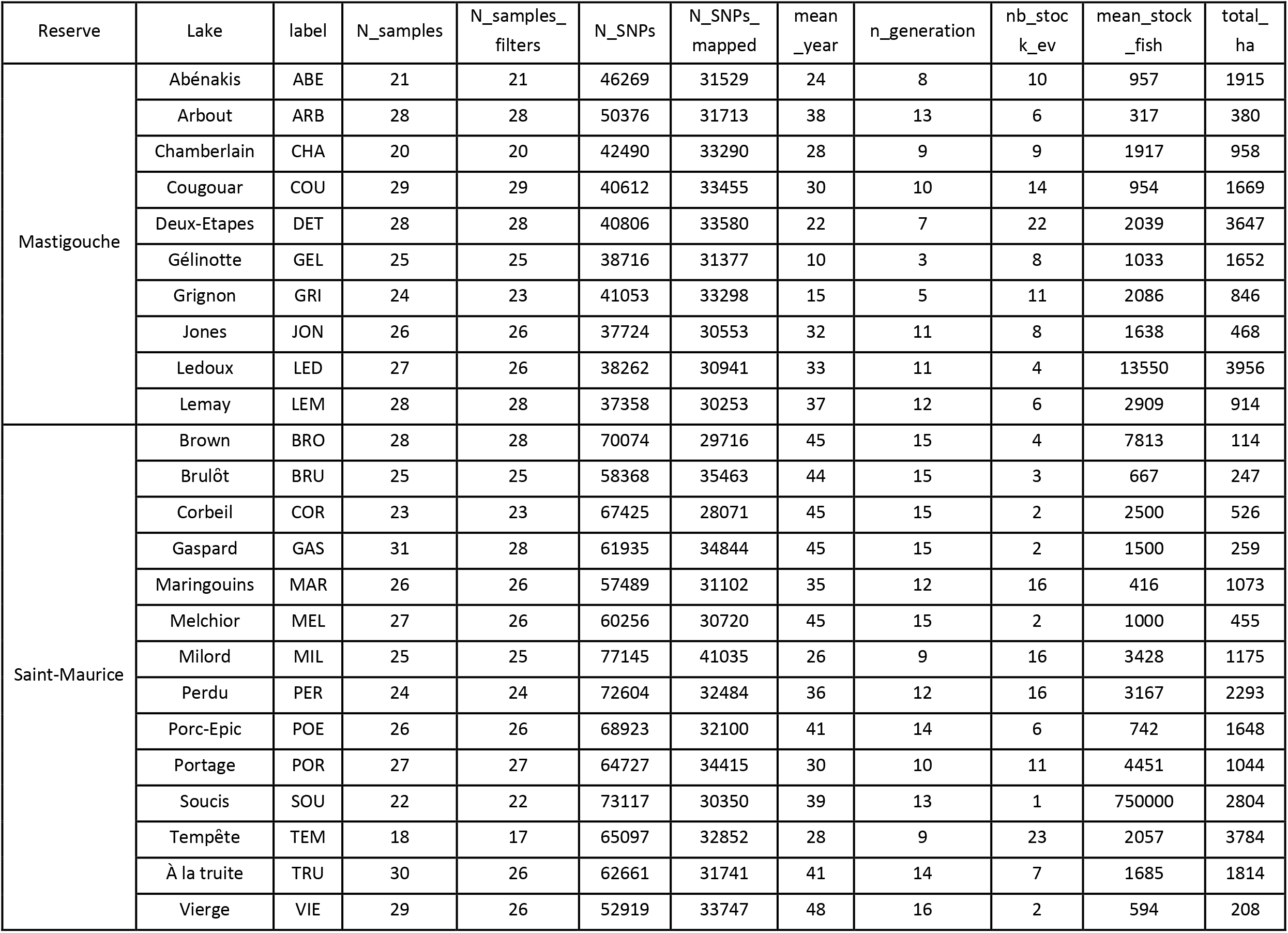
Description of the 24 sampled Brook Charr lakes in Québec, Canada along with the stocking variables: N_samples: number of individuals per lake; N_samples_filters: number of individuals after filtering; N_SNPs: the number of SNPs after filtering; N_SNPs_mapped: the number of mapped SNPs; mean_year: the number of years since the mean year of stocking; nb_stock_ev: the total number of stocking events; mean_stock_fish: the mean number of fish stocked per stocking event and total_ha: the total number of fish stocked per stocking event.

| Reserve | Lake | label | N_samples | N_samples_filters | N_SNPs | N_SNPs_mapped | mean_year | n_generation | nb_stock_ev | mean_stock_fish | total_ha |
| --- | --- | --- | --- | --- | --- | --- | --- | --- | --- | --- | --- |
| Mastigouche | Abénakis | ABE | 21 | 21 | 46269 | 31529 | 24 | 8 | 10 | 957 | 1915 |
|  | Arbout | ARB | 28 | 28 | 50376 | 31713 | 38 | 13 | 6 | 317 | 380 |
|  | Chamberlain | CHA | 20 | 20 | 42490 | 33290 | 28 | 9 | 9 | 1917 | 958 |
|  | Cougouar | COU | 29 | 29 | 40612 | 33455 | 30 | 10 | 14 | 954 | 1669 |
|  | Deux-Etapes | DET | 28 | 28 | 40806 | 33580 | 22 | 7 | 22 | 2039 | 3647 |
|  | Gélinotte | GEL | 25 | 25 | 38716 | 31377 | 10 | 3 | 8 | 1033 | 1652 |
|  | Grignon | GRI | 24 | 23 | 41053 | 33298 | 15 | 5 | 11 | 2086 | 846 |
|  | Jones | JON | 26 | 26 | 37724 | 30553 | 32 | 11 | 8 | 1638 | 468 |
|  | Ledoux | LED | 27 | 26 | 38262 | 30941 | 33 | 11 | 4 | 13550 | 3956 |
|  | Lemay | LEM | 28 | 28 | 37358 | 30253 | 37 | 12 | 6 | 2909 | 914 |
| Saint-Maurice | Brown | BRO | 28 | 28 | 70074 | 29716 | 45 | 15 | 4 | 7813 | 114 |
|  | Brulôt | BRU | 25 | 25 | 58368 | 35463 | 44 | 15 | 3 | 667 | 247 |
|  | Corbeil | COR | 23 | 23 | 67425 | 28071 | 45 | 15 | 2 | 2500 | 526 |
|  | Gaspard | GAS | 31 | 28 | 61935 | 34844 | 45 | 15 | 2 | 1500 | 259 |
|  | Maringouins | MAR | 26 | 26 | 57489 | 31102 | 35 | 12 | 16 | 416 | 1073 |
|  | Melchior | MEL | 27 | 26 | 60256 | 30720 | 45 | 15 | 2 | 1000 | 455 |
|  | Milord | MIL | 25 | 25 | 77145 | 41035 | 26 | 9 | 16 | 3428 | 1175 |
|  | Perdu | PER | 24 | 24 | 72604 | 32484 | 36 | 12 | 16 | 3167 | 2293 |
|  | Porc-Epic | POE | 26 | 26 | 68923 | 32100 | 41 | 14 | 6 | 742 | 1648 |
|  | Portage | POR | 27 | 27 | 64727 | 34415 | 30 | 10 | 11 | 4451 | 1044 |
|  | Soucis | SOU | 22 | 22 | 73117 | 30350 | 39 | 13 | 1 | 750000 | 2804 |
|  | Tempête | TEM | 18 | 17 | 65097 | 32852 | 28 | 9 | 23 | 2057 | 3784 |
|  | À la truite | TRU | 30 | 26 | 62661 | 31741 | 41 | 14 | 7 | 1685 | 1814 |
|  | Vierge | VIE | 29 | 26 | 52919 | 33747 | 48 | 16 | 2 | 594 | 208 |

### Inference of local ancestry

Local ancestry inference was performed following the same methodology developed by Leitwein et al. (2018). First, we used the program ELAI v1.01 (Guan, 2014) based on a two-layer hidden Markov model to detect individual ancestry dosage from the domestic strain along each individual linkage group (LG hereafter). The program was run 20 times for each 42 Brook Charr LGs to assess convergence. Prior to running ELAI, we retrieved the relative mapping positions of our makers along each chromosome after controlling for synteny and collinearity between the Arctic Charr and the Brook Charr genomes. To identify blocks of conserved synteny between the two species, we anchored both the Brook Charr and the Artcic Charr linkage maps (Sutherland et al., 2016 and Nugent, Easton, Norman, Ferguson, & Danzmann, 2017, respectively) to the Arctic Charr reference genome using MAPCOMP (Sutherland et al., 2016). Results were visualized with the web-based VGSC (Vector Graph toolkit of genome Synteny and Collinearity: http://bio.njfu.edu.cn/vgsc-web/). We were thus able to order RAD loci (that were assembled against the Arctic Charr reference genome) with respect to their relative positions along each of the 42 the Brook Charr LGs before running ELAI. We then performed local ancestry inference separately for each population, using the 33 domestic individuals as a source population and the wild caught individuals as the admixed population. For each LG in each of the 24 populations, 20 replicate runs of ELAI were performed with the number of upper clusters (-C) set to 2 (i.e. assuming that each fish was a mixture of domestic and wild populations), the number of lower clusters (-c) to 15, and the number of expectation-maximization steps (-s) to 20. Finally, the number of admixture generations (-mg) was estimated using the mean year of stocking and the approximate mean age at maturity of 3 years and ranged from 3 to 16 generations (Table 1). We then generated individual domestic ancestry profiles by plotting the estimated number of domestic allele copies for each replicate run and its median along each LG with R (Team, 2015). The pipeline we used is available on GitHub (https://github.com/mleitwein/local_ancestry_inference_with_ELAI). We then compared the percentage of domestic ancestry computed here with ELAI to the previous study from Létourneau et al. (2018) using a spearman’s correlation.

### Estimation of domestic ancestry tracts length and number

The number and length of domestic tracts were determined based on the positions of junctions retrieved from ELAI ancestry dosage output, following the method used and detailed in Leitwein et al. (2018). When the ELAI domestic ancestry dosage median value was comprised between [0.9 to 1.1] and [1.9 to 2], we considered the presence of one domestic tracts (i.e. in heterozygous state), and two domestic tracts (i.e. in homozygous state), respectively. Junction positions within “uncertainty areas” (when the domestic ancestry dosage was either comprised between 0.1 and 0.9 or between 1.1 and 1.9) were determined as the position where the domestic ancestry dosage crossed the 0.5 or 1.5 value (see Leitwein et al. (2018) for details). Junction positions were used to estimate the number and length of introgressed domestic tracts for each of the 42 LGs in each population.

### Hybrids class determination

In order to define hybrid categories with respect to the number of generations of crossing in nature, we computed the Chromosomal Ancestry Imbalance (CAI) developed by Leitwein et al., (2018). Briefly, the CAI represents the cumulated length differences of the domestic ancestry between the two parental chromosome copies divided by the respective linkage group length. Thus, pure domestic or pure wild individuals have a CAI of 0 whereas F1 hybrids between wild and domestic parents have a CAI of 1. Due to uncertainty concerning individual haplotype structure (i.e. local ancestry profiles were inferred from unphased domestic ancestry dosage), we could not precisely determine the CAI of admixed genotypes resulting from several generations of admixture. Therefore, we conservatively classified individuals as early-generation hybrids when their CAI was equal or higher than 0.25, and as late-generation hybrids when their CAI was equal or lower than 0.125. Early generation hybrids therefore correspond to F1, F2, first generation backcrosses and other types of crosses generated among hybrids and parental pedigrees during the first generations of admixture. By contrast, late generation hybrids comprise genotypes that are mostly made of wild-type ancestry, while being introgressed by varied proportions of domestic alleles. In both hybrid categories, we removed individuals with a genome-wide percentage of domestic ancestry higher than 60% (see Figure Sup 1) in order to exclude individuals with a mostly domestic ancestry. This won’t impact the following analysis as these individuals are probably pure domestic individuals or hybrids between F1 and domestic parents; this concerns only six individuals (Figure Sup1).

### Impact of stocking on domestic haplotype number and length

We first assessed the relationships between domestic tract characteristics and the four variables reflecting the stocking history (i.e., the mean_year, the nb_stock_ev, the mean_stock_fish and the total_ha; Table 1) using linear mixed models. The mean percentage of domestic ancestry, the number and length of introgressed domestic tracts were treated as dependent variable whereas the stocking variables were introduced in the model as explanatory terms in an additive way, with no interaction to avoid model over-parameterization. The dependent variables were log-transformed and the explanatory variables were scaled (i.e., centered and reduced). The population (24 populations) and the region (two regions, Mastigouche and StMaurice) were introduced as random effects in the model. Normality of the residuals was examined graphically using a quantile–quantile plot. We used a likelihood ratio test to assess the significance of the tested relationship by comparing the models with and without the explanatory term. We calculated marginal R^2^ to quantify the proportion of variance explained by the explanatory variable only. All analyses were performed in R using the package “lme4” (Bates, Mächler, Bolker, & Walker, 2015).

### Domestic ancestry profiles as a function of hybrid classes

The fraction of domestic ancestry rate was estimated separately for both hybrid classes, with the early-generation hybrids comprising 47 individuals from 17 populations and the late-generation hybrids comprising 394 individuals from the 24 populations. Local ancestry rate from the domestic strain was estimated at 11,803 SNPs positions distributed along the 42 LGs and common between all the individuals originating from different lakes and found in the two hybrids categories. Estimates of genome-wide domestic ancestry rate for each hybrid category were plotted with R (Team, 2015) (https://github.com/mleitwein/local_ancestry_inference_with_ELAI). The mean domestic ancestry rate and its 95% CI were reported along the 42 LGs. Then, genomic regions exceeding the 95% CI were considered as displaying excess or deficit of domestic ancestry.

Linear mixed models were used to investigate how domestic ancestry profiles (i.e. mean number and length of domestic tracts) differ between early and late hybrid classes. The domestic tracts characteristics were log-transformed and treated as dependent variables whereas the hybrid class (discrete variable with two modalities) was introduced in the model as an explanatory term. The population (24 populations) and region (two regions, Mastigouche and St Maurice) terms were introduced as random effects in the model. All analyses were performed in R using the package “lme4” (Bates et al., 2015).

### Local recombination rate estimation

In order to estimate genome-wide variation in recombination rate, we anchored the Brook Charr mapped RAD loci from Sutherland et al. (2016) to the Arctic Charr reference genome. The relative position of these loci were extracted from the BWA_mem alignment for each collinearity block identified with MAPCOMP (Sutherland et al., 2016) allowing the reconstruction of a Brook Charr collinear reference genome. Then, the local variation in recombination rate was estimated across the collinear reference genome by comparing the physical (bp) and genetic position (cM) of each marker using MareyMap (Rezvoy, Charif, Guéguen, & Marais, 2007). The polynomial Loess regression method was used to assess the recombination rate with a degree of smoothing (*span*) set to 0.9. Finally, to estimate the recombination rate of markers that were not included in the linkage map, we computed the weighted mean recombination rate using the two closest markers based on their relative physical positions.

### Relationships between domestic ancestry and recombination rate

To investigate the selective forces shaping the domestic ancestry pattern along the genome, we took in consideration the recombination rate. Indeed, a positive correlation between recombination rate and domestic ancestry rate reflect selective forces against domestic ancestry in low recombining regions. Inversely, a negative correlation will reflect selective forces favoring domestic ancestry in low recombining regions. As the recombination rate is highly variable along the genome, we investigated the relationship between the local domestic ancestry rate and the recombination rate at three different levels: (i) at the whole genome level, (ii) at the linkage group level and (iii) within 2Mb windows. For the three levels, we used regression models in which the local domestic ancestry rate was treated as dependent variable and the recombination rate was incorporated as an explanatory term. In all models, domestic ancestry rate was log-transformed and the recombination rate was centered-reduced. At the whole genome level, we used a linear mixed model in which the linkage groups and the 2Mb windows were introduced as random effects. By doing so, we considered the non-independency of the domestic ancestry rate estimates by specifying that the estimates belong to a given window within a given linkage group. The hybrid class (early and late) was also added as an explanatory term in an interactive way (recombination × class). The marginal R^2^ describing the proportion of variance explained by the fixed factors (i.e. the recombination rate and the hybrids class) was calculated and we assessed the significance of the hybrid class effect using a likelihood ratio test. At the linkage group level, in order to avoid model over-parameterization, the analyses were performed separately for the early and late hybrids. A linear mixed model was built for each of the 42 linkage groups and the 2Mb sliding window was introduced as a random effect in all models. The slope coefficient (and the 95% CI) of the relationship between domestic ancestry rate and recombination rate was reported for the 42 linkage groups. At the 2Mb window level, generalized linear models were built for each window containing at least 5 positions with an estimated recombination and introgression rate (resulting in 689 models) because a regression analysis could not be performed with fewer values. The slope coefficient and its 95% CI were reported for each sliding window. In all models, the significance of the effect of the explanatory terms was assessed with a likelihood ratio test. All analyses were performed in R using the package “lme4” (Team, 2015).

### Relationships between potentially deleterious mutations and ancestry rate

All SNPs present in all populations were used for the identification of potentially deleterious mutations. First, read sequences associated to the SNPs were blasted against the Arctic Charr proteome available on NCBI (https://www.ncbi.nlm.nih.gov/assembly/GCF_002910315.2). All hits with a minimum amino acid sequence length alignment of 25 and a similarity of 70% between the sequences of interest and the reference proteome were retained (those thresholds have been optimally chosen after running several tests with different parameter combinations on the observed data). Then, PROVEAN (Protein Variation Effect Analyzer; Choi, Sims, Murphy, Miller, & Chan, 2012) was used to predict the deleterious effect of nonsynonymous mutations. As in previous studies (e.g. Renaut & Rieseberg, 2015; Ferchaud, Laporte, Perrier, & Bernatchez, 2018) a threshold of −2.5 in Provean score was applied to distinguish between nonsynonymous mutations potentially deleterious (≤-2.5) and neutral (>-2.5). In PROVEAN, the deleteriousness of a variant can be predicted based on its effect on gene functioning (such as protein changing, stop-gain, stop-lost), for example by assessing the degree of conservation of an amino acid residue across species. The pipeline used for the entire process is available on github (gbs_synonymy_genome: https://github.com/QuentinRougemont/gbs_synonymy_with_genome).

To evaluate the relationship between the presence of potentially deleterious mutations and the domestic ancestry rate as a function of recombination rate, a mean domestic ancestry and recombination rate was estimated in a window size of 400Kb surrounding the position of the potentially deleterious mutation (i.e. 200Kb before and after). The analysis was performed at the genome-wide level and linkage group level. For both levels, we first applied a regression model in which the mean 400Kb window domestic ancestry rate was treated as a dependent variable and the associated recombination rate window was incorporated as an explanatory term. At the genome-wide scale, a linear mixed model was performed and linkage groups were incorporated as random effect and the recombination rate as a control co-variable. We then performed a linear model in which the residuals of the model (domestic ancestry rate ~ recombination rate) were included as a dependent variable and the presence of potentially deleterious alleles (Provean score<-2.5) or neutral (Provean score >-2.5) as an explanatory term (i.e. discrete variable with two modalities, “deleterious” vs “neutral”). At the linkage group level, similar linear models were built for each linkage group.

## Results

### SNPs calling

Demultiplexed and cleaned raw reads resulted in an average of 2.38 million reads per individual. After filtering, 636 individuals were kept for further analyses. On average, 1,225,704 ± 285,583 reads were properly mapped to the Arctic Charr reference genome with an average depth of 11.5X ± 2.4 per individual. After applying population filters, an average of 55,267 ± 11,779 SNPs were kept per population for subsequent analysis (Table 1).

### Domestic ancestry tracts detection and hybrid classes

We performed local ancestry inference with ELAI with an average of 32,442 ± 1,799 SNPs per population that were mapped to the reconstructed Brook Charr reference genome (see methods and Table1). Individual local ancestry was summarized across linkage groups to determine the number and mean length of domestic ancestry tracts at the individual level. Both the domestic ancestry tract length and abundance were highly variable among individuals, ranging from 356Mb to 72 Mb in length and from 0 to 103 for the number of domestic ancestry tracts per individual (Table Sup. 1 and Table Sup. 2). A total of 47 individuals were assigned to the early hybrid-generation class (CAI>=0.25) and 394 to the late hybrid-generation class (CAI<=0.125) (Table Sup. 1). A total of 195 remaining fish where unclassified and therefore considered as intermediate between early and late-generation hybrid categories based on these empirical thresholds. The mean individual percentage of domestic ancestry was significantly positively correlated (Spearman’s rho = 0.68, P < 10^−10^, Figure Sup 2) with the percentage of domestic ancestry computed with ADMIXTURE by Létourneau et al., (2018) but that was based on a much smaller number of markers (~4,579 SNPs per populations).

### Influence of stocking history on domestic tract number and length

No significant relationship was found between the mean number of years since stocking or the intensity of stocking and the mean individual percentage of domestic ancestry within each population (Table Sup. 1). However, the number of domestic tracts tended to be higher for populations that have been supplemented enhanced more than 30 years ago. Indeed, a marginally significant positive relationship was detected between the mean number of domestic tracts and the number of years since the mean year of stocking (R^2^=0.08, p-value=0.06, Figure 1 A). Also, a significant negative correlation was observed between the mean length of individual domestic tracts and the number of years since the mean year of stocking (Figure 1 B, R^2^ =0.055, p-value <0.01). Therefore, the length of the domestic ancestry tracts tended to decrease over time, whereas it tended to increase with the number of stocking events (Figure Sup 3, R^2^=0.055, p-value= 0.049).

**Figure 1.**
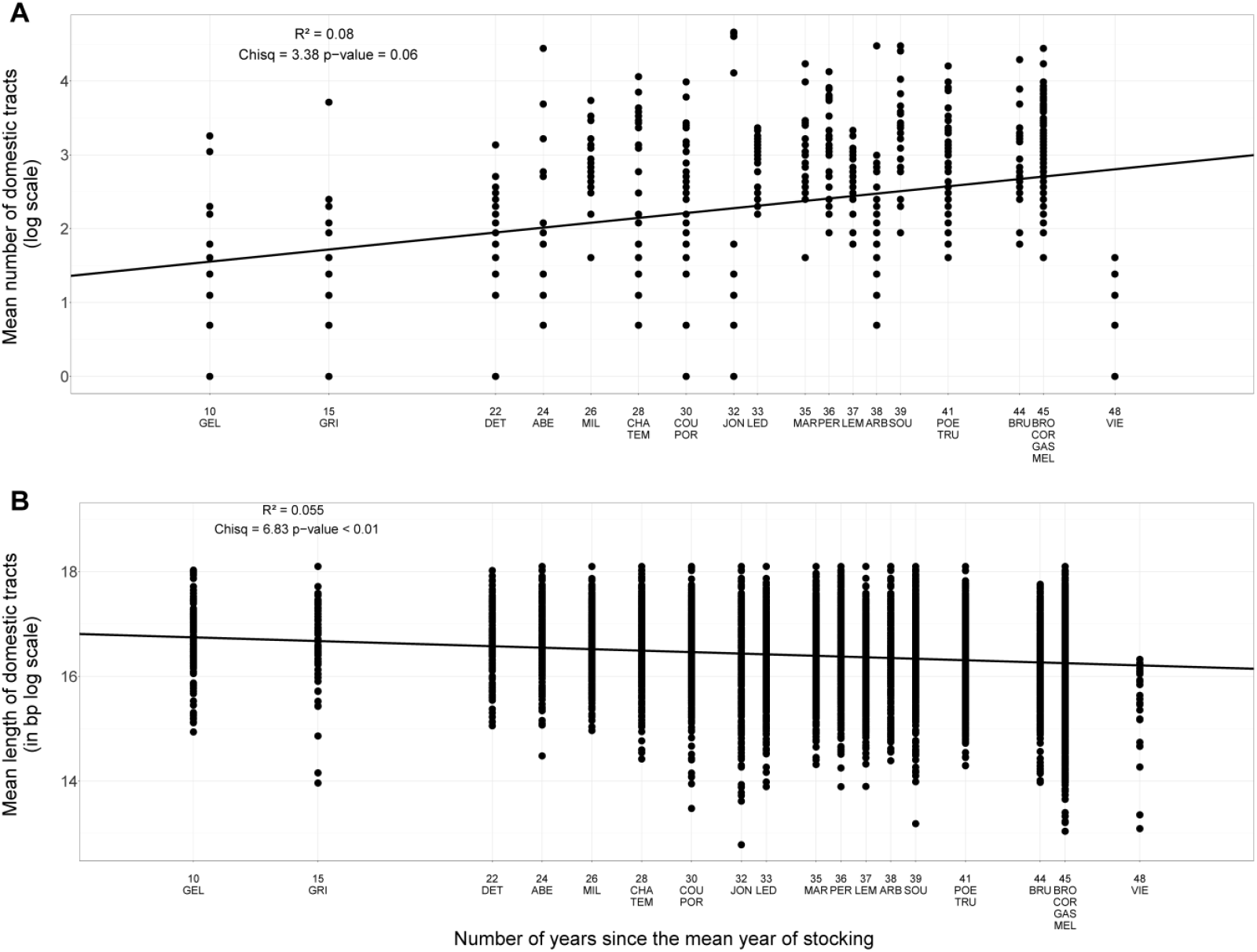
Relationships between domestic tract features (mean number and length) and stocking variables. A: Positive correlation between the mean number of domestic tracts (log scale) per individual as a function of the number of years since the mean year of stocking events for each sampled lake (lakes labels are described in table 1; R^2^=0.08, p-value=0.06). B: Negative correlation between the mean length of domestic tracts in bp log scale per individual as a function of the number of years since the mean year of stocking events for each sampled lake (lakes labels are described in table 1; R^2^=0.055, p-value<0.01).

### Domestic tracts characteristics as a function of hybrid classes

The mean number of domestic ancestry tracts was significantly lower for the late hybrid individuals (*mean*_*number*_= 8.01) compared to the early hybrid individuals (*mean*_*number*_ = 46.31) (Figure 2 A, R^2^=0.22, p-value <2.2e-16). Moreover, the mean length of the introgressed domestic tracts was shorter for the late hybrid (*mean*_*length*_= 14,6 Mb) compared to early hybrid individuals (*mean*_*length*_=19,7 Mb bp) (Figure 2 B, R^2^=0.06, p-value < 3.9e-10).

**Figure 2.**
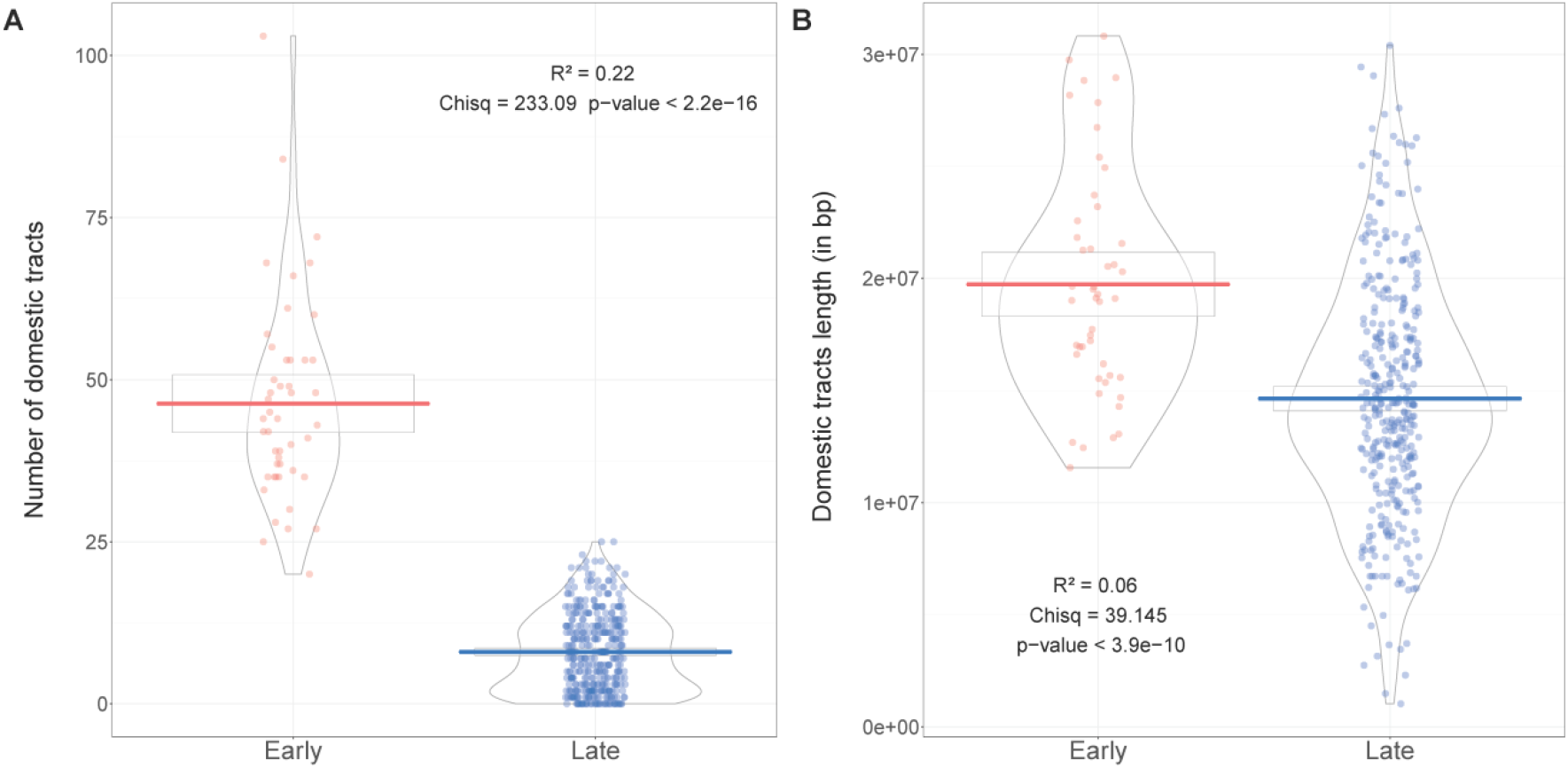
Domestic tract features and hybrid generations (early and late). The number (A) and the length (in bp) (B) of domestic tracts as a function of the hybrid categories; early (in pink) and late (in blue). Boxes indicate 95% CI, horizontal line represents the mean and violins indicate the density

### Domestic ancestry and recombination rates

The genome-wide level of domestic ancestry rate was estimated for both early and late hybrid-generations using 11,803 SNPs (i.e. corresponding to common polymorphic sites among the two hybrid categories) distributed along the 42 Brook Charr linkage groups. The mean genome-wide domestic ancestry rate in early hybrid-generations was 0.307 (95% CI 0.18-0.43) (Figure Sup. 4A) and was highly variable within and among linkage groups with several genomic regions displaying a local excess (LG1, 9, 11 and 12, Figure Sup. 4A) or deficit of domestic ancestry (LGs 14, 28, 30 and 40; Figure Sup. 4A). The mean domestic ancestry rate of late hybrid-generations was significantly lower with a mean of 0.042 (95% CI 0.012-0.081) (Figure Sup. 4A), and genomic regions displaying excess (LGs 9, 12, 20 and 41) or deficit (LG 14) of domestic ancestry were found (Figure Sup. 4A). Thus, two linkage groups displaying an excess (LGs 9 and 12) and one displaying a local deficit of domestic ancestry (LG 14) were found both in early and late hybrid-generations.

The local recombination rate (r) was estimated at 10,740 SNPs out of the 11,803 SNPs and was found to be highly variable within and among linkage groups, with a mean value of 4.48 cM/Mb ± 1.83 (Figure Sup. 4B) over the 42 linkage groups. At the genome-wide scale, for both early and late-generation hybrids, a negative correlation was found between the proportion of domestic ancestry and recombination rate (Figure 3A; slope coefficient β_**early**_= −0.07,β _**Late**_ =-0.09; p-value<2.2e-16). The marginal R^2^ was equal to 0.90, the likelihood ratio test was significant for the recombination rate (χ^2^ _Recombination_= 441.7, p-value<2.2e-16), the hybrid class (χ^2^ _Hybrids_= 60420, p-value<2.2e-16) and the interaction between recombination and the hybrid class (χ^2^ _Hybrids_interaction_=177.57; p-value<2.2e-16). Moreover, the interaction between recombination rate and hybrid class was significant (β_**early:Late**_ =2.8; p-value<2.2e-16), indicating a stronger negative correlation between introgression and recombination rate in late compared to early-generation hybrids.

**Figure 3.**
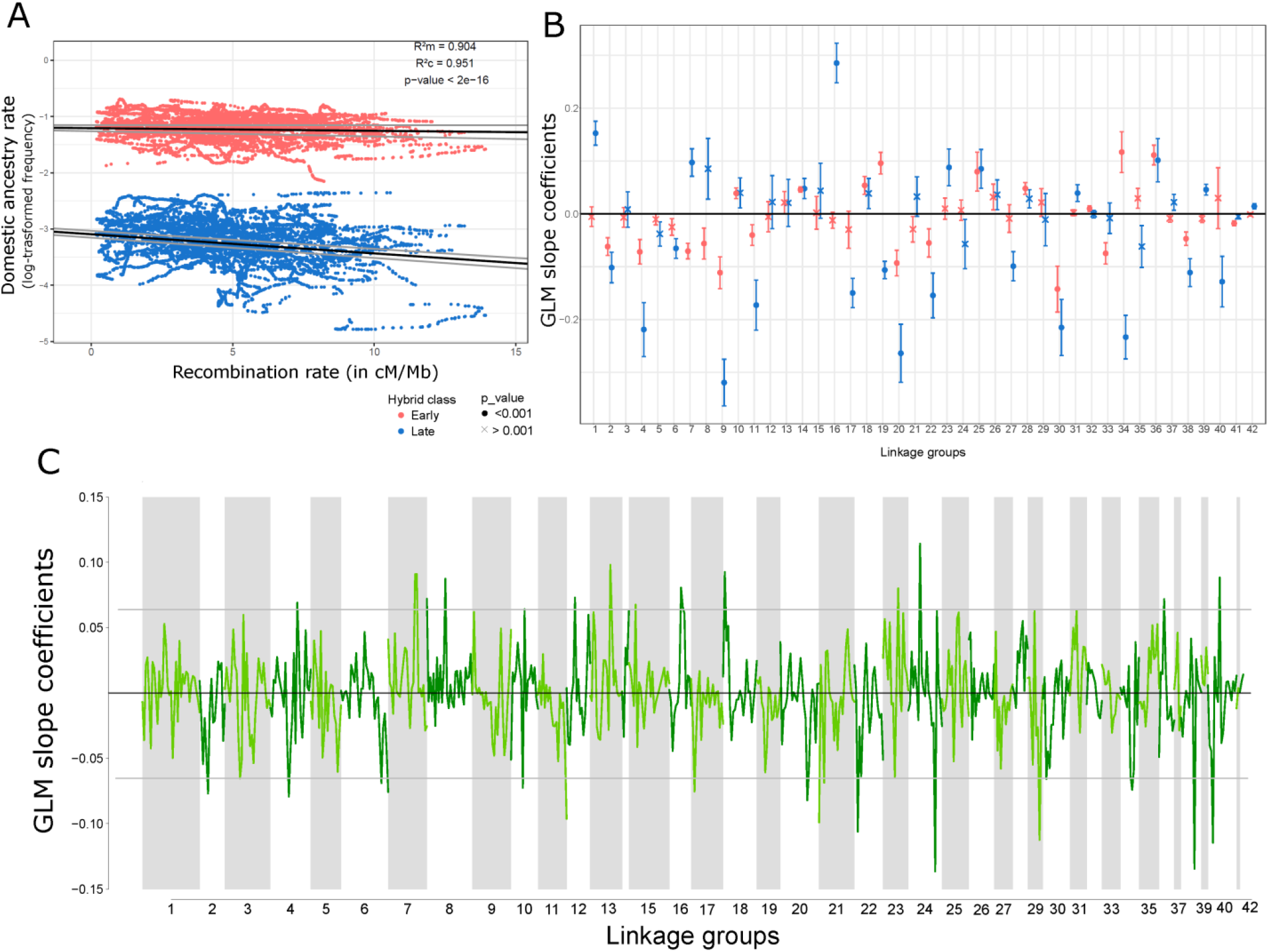
Relationship between the domestic introgression rate and the recombination rate in cM/Mb at different genomic scales. A) The whole genome scale; negative correlation between the log-transformed introgression rate (frequency among individuals) and the recombination rate (cM/Mb) for both early (in pink) and late (in blue) hybrids categories assessed with a generalized linear mixed models (GLM; *β*_early_ = −0.07, *β*_Late_=-0.09; R^2^m=0.90, R^2^c=0.95; LRtest χ^2^ _Recombination_= 441.7, χ^2^ _Hybrids_= 60420, χ^2^ _Hybrids_interaction_=177.57; p-value<2.2e-16). B) The linkage group level; slope coefficient of the GLM of the introgression rate and recombination rate models for each 42 Brook Charr linkage group for both early (in pink) and late (in blue) hybrid categories. The dot shapes represent the significance of each model with the dark circle and cross representing p-value<0.001 and p-value>0.001, respectively. C) The 2Mb sliding windows level; slope coefficient of the GLM of the introgression rate and recombination rate models of the late hybrids for each 2Mb sliding windows along the 42 Brook Charr linkage groups. Grey lines represent the 95% confidence interval.

The results of the linear mixed model fitted at the linkage group level are presented in Figure 3B (mean and 95% CI) for both hybrids classes. For the late-generation hybrids, 10 linkage groups showed a significant positive correlation and 14 linkage groups showed a significant negative correlation between the domestic ancestry rate and recombination rate (Figure 3B). For the early-generation hybrids, nine and 12 linkage groups respectively showed a significant positive and a significant negative correlation between the domestic ancestry rate and recombination rate (Figure 3B). Eight linkage groups displayed significant negative correlations between the domestic ancestry and recombination for both early and late-generation hybrid classes, whereas only 3 LG consistently showed positive correlations. Conversely, three LGs presented opposite correlations between the early and the late-generation hybrid classes (LGs 7, 19 and 34).

We then focused the window-scale analyses on late-generation hybrids only because the effect of selection needs more than a few generations after hybridization to leave detectable footprints at such a local scale. The estimated slopes of linear models performed for each 2Mb sliding windows in late-generation hybrids were highly variable within and among linkage groups, and several genomic regions exceeded the 95% confidence interval of the estimated distribution (grey lines; Figure 3C). For example, 12 genomic regions showed a strong positive (e.g. LGs 7, 8, 13, 16, 24, 40) correlation and 15 genomic regions showed a strong negative correlation (e.g. LGs 2, 12, 22, 24, 29, 38, 40; Figure 3C) between local domestic ancestry rate and recombination rate.

### Potentially deleterious mutations

Among a total of 141,332 sequences that were blasted against the Arctic Charr reference proteome (Christensen et al., 2018), 36,743 loci had significant hits and were retrieved. Among these, 21,288 had more than 70% similarity between the sequences of interest and the reference proteome and a mean length greater than 25 nucleotides. Finally, 7,799 were non-synonymous and among them 1,932 were potentially deleterious (PROVEAN score <-2.5).

At the genome scale, we did not detect any significant correlation between the residuals of the linear relationship between local ancestry and recombination rate and the presence of deleterious mutations, in both early (β_**early**_= −0.0015; p-value= 0.6137) and late-generation hybrids (β_**Late**_=-0.0114; p-value=0.0713) (Figure Sup. 5). At the linkage group scale, two linkage groups (LGs 4 and 15) showed a negative relationship between the residuals and the presence of potentially deleterious alleles for the early hybrids (Figure Sup. 5A and B; LG4: R^2^=0.018, p-value=0.04; LG15: R^2^=0.02, p-value=0.028). For the late hybrids, two linkage groups showed a significant negative correlation, LG7 (R^2^=0.02, p-value= 0.045) and LG41 (R^2^=0.20, p-value=0.01), and two linkage groups showed a marginally significant positive correlation, LG20 (R^2^=0.01, p-value=0.059) and LG26 (R^2^=0.033, p-value=0.051), between the presence of potentially deleterious alleles and residuals of the linear mixed model of domestic ancestry rate and the recombination rate (Figure 4).

**Figure 4.**
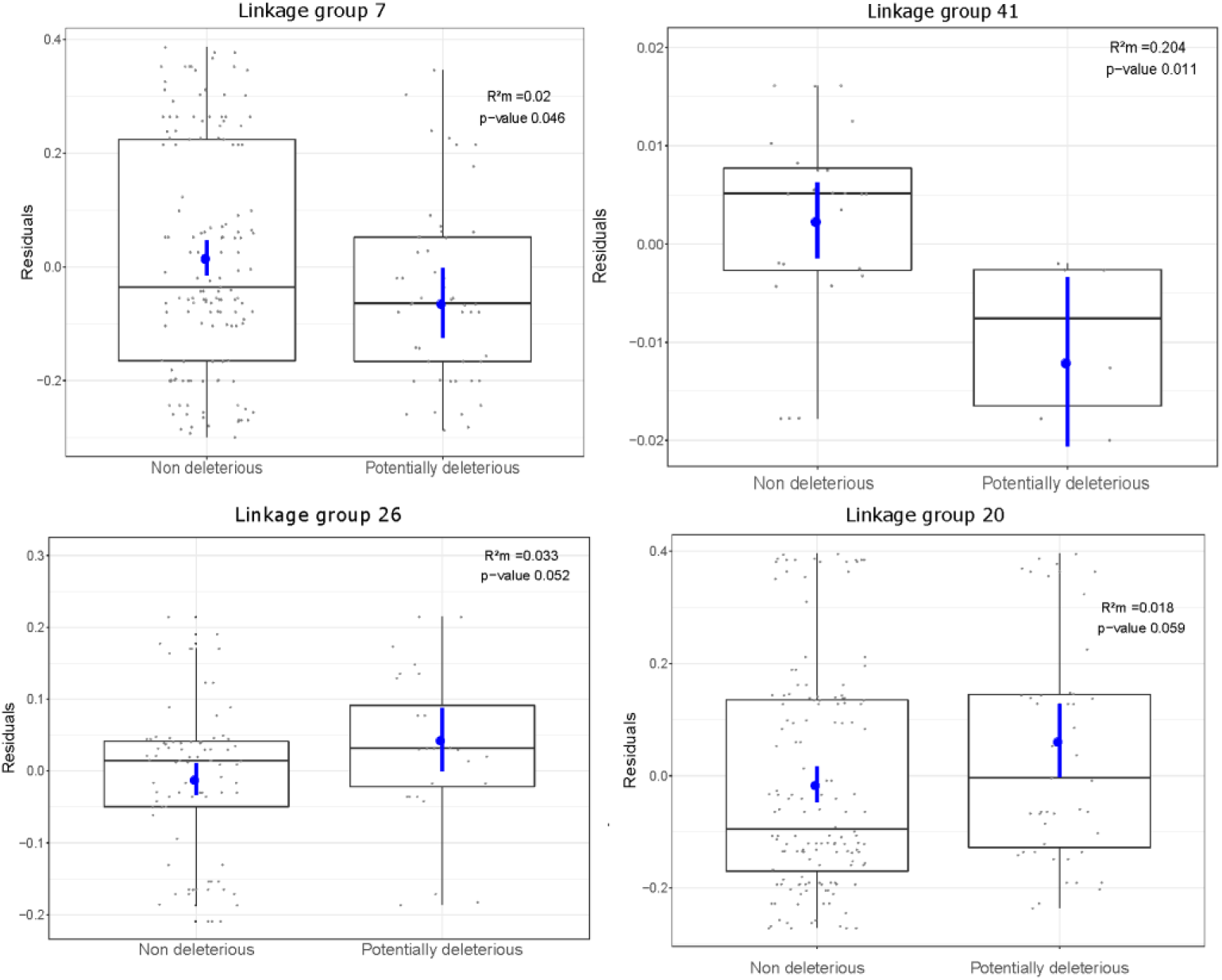
Domestic ancestry rate excess or deficit and presence of potentially deleterious alleles. Residuals of the regression between the domestic ancestry and recombination rates models as a function of the presence of non-deleterious or potentially deleterious alleles for four linkage groups in late hybrids generations. LG7 and LG41 displayed a lower introgression rate of domestic ancestry compared to model prediction in genomic regions surrounding potentially deleterious alleles. LG26 and LG20 displayed an excess of introgressed domestic ancestry compared to the models expectation around potentially deleterious alleles. Boxes indicate 95% confidence intervals, horizontal line represents the median. Blue dots and blue line represent the estimates and the confidence intervals of the linear model estimates. Marginal R^2^ (R^2^m) and significance are displayed at the upper right of each plot.

## Discussion

The main goal of this study was to document the genetic outcomes of more than four decades of stocking domestic Brook Charr into wild populations. To this end, we assessed genome-wide patterns of domestic ancestry within 24 wild populations with different histories of stocking. We more specifically considered the relationship between recombination rate and domestic ancestry rate at different scales: (i) the whole genome, (ii) individual linkage groups and (iii) within 2Mb windows. This allowed us to detect a wide range of patterns that can be attributed to different evolutionary processes acting across the genome. Among these, the main factor modulating local domestic ancestry was likely associative overdominance leading an increased frequency of domestic ancestry within low recombining regions. This pattern might be explained by the presence of mildly deleterious recessive mutations in the wild populations. At the regional scale (i.e. chromosomes and 2Mb windows), inversed correlations were found which suggested local selection against domestic ancestry. Our results highlight the importance of taking into consideration genome-wide variation in both recombination and ancestry rates to understand the evolutionary outcomes of human induced admixture.

### The history of supplementation impacts domestic ancestry

No significant relationship was observed between the mean individual proportion of domestic ancestry and the main stocking variables previously identified as having an impact on the levels of domestic introgression (i.e. mean year since stocking, number of stocking event, number of fish stocked per ha and the number of stocking events; Létourneau et al., 2018). This discrepancy could be explained by different sensitivities of the methods. Indeed, although the correlation between individual domestic ancestry proportions estimated here and in Létourneau et al. (2018) was good (Spearman’s rho = 0.68, P < 10^−10^), low proportions of domestic ancestry were not detected in the previous study, which resulted in considering weakly introgressed individuals as pure wild genotypes (Figure Sup 2). Moreover, thanks to the higher number of markers and their positions we were able to retrieve the approximated length and number of domestic tracts within each population. Consistent with theoretical predictions (Racimo et al., 2015), we observed an increase in the number of domestic ancestry haplotypes and a decrease in the mean domestic haplotype length as a function of the number of years since the main stocking event. This corroborates earlier findings by Leitwein et al. (2018) who suggested that the mean length of foreign ancestry could be used as a proxy to retrieve the history of stocking practice. We indeed observed a higher number of smaller introgressed domestic haplotypes for the lakes where stocking has stopped earlier in the past. Additionally, the length and the number of domestic tracts also allowed distinguishing among early and late-generation-hybrids within populations, which is important as the lakes have undergone several successive events of supplementation. Considering the time since hybridization by distinguishing early (i.e. F1, F2, and backcrosses of first hybrid generations) and late-generation hybrids (i.e. individuals with small number of short domestic haplotypes) is also important to assess the potential evolutionary outcomes of supplementation. Indeed, selection is expected to be more efficient in later hybrid generations when introduced domestic haplotypes have been sufficiently shortened by recombination.

### General tendency of selective effects and putative deleterious mutations

Both the time since hybridization and the genomic scales (i.e. global *versus* local scales) were important to assess the selective outcomes of gene flow between wild and domestic individuals. At the genome-wide scale, we observed a negative correlation between domestic ancestry and recombination rate for both first and late-generation hybrids. While we detected highly variable selective effects along the genome, this negative correlation was also predominant at the linkage group and at the local scales (i.e. 2Mb windows) for the late hybrid generations. Such negative correlation could be the result of different evolutionary mechanisms. First, the presence of beneficial mutations under positive selection which may locally increase the introgression of foreign alleles in low recombining region where hitchhiking of neutral foreign alleles could occur (Felsenstein, 1974; B. Charlesworth, 2009; Fay & Wu, 2000). For example, adaptive introgression of Neanderthal ancestry has been reported in modern humans and interpreted as an adaptation to high altitude in Tibetan populations (Huerta-Sánchez et al., 2014) and the immune response (Dannemann & Racimo, 2018; Racimo et al., 2015). However, this mechanism is more likely to explain local scale correlations than genome-wide patterns. Secondly, given the generally small effective population size of these Brook Charr lacustrine populations (Gossieux, Bernatchez, Sirois, & Garant, 2019), random drift may also be responsible for variable ancestry locally along the genome (S. Martin & Jiggins, 2017) and is not expected to produce consistent patterns across lakes. Thirdly, the most likely evolutionary mechanism that could explain the general negative correlation would be a dominant effect of associative overdominance (Kim et al., 2018). Indeed, favored domestic ancestry within low recombination rate regions is likely to reflect the action of associative-overdominance, especially if the recipient population tends to accumulate recessive deleterious alleles mostly in low recombining regions (D. Charlesworth & Willis, 2009). Consequently, long introgressed foreign haplotypes of domestic origin would be favored by masking the effect of linked recessive deleterious mutations present in the small local populations (i.e. local heterosis effect; Charlesworth & Willis, 2009; Kim et al., 2018). Kim et al. (2018) suggested that the presence of slightly deleterious recessive mutations might modulate the genome-wide introgression rate in hybrid populations. More specifically, if the mutation load of receiving (wild) populations is higher than in the introduced domestic strain, a negative correlation between domestic ancestry rate and recombination rate is expected (Schumer et al., 2018). Here, when considering the genome-wide scale, we did not observe any relationship between the presence of putative deleterious mutations and the residuals of the linear model relating introgression to recombination rate. However, for some of the chromosomes (LGs 26 and 20), we were able to detect an increase of domestic ancestry in the presence of deleterious mutations which may reflect a positive effect (i.e. associative over-dominance) of the presence of domestic ancestry. This is congruent with the hypothesis of temporary reduced genetic load, caused by the masking of deleterious mutations, and resulting in an increase of introgressed ancestry as observed in Kim et al. (2018). This is also consistent with the expected higher accumulation of deleterious mutations in small lacustrine Brook charr populations as observed in Ferchaud et al. (2019) as well as with the higher allelic richness observed in the domestic strain compared to these small lacustrine populations (S. Martin, Savaria, Audet, & Bernatchez, 1997).

### Variability of selective effects revealed at local scales

At local genomic scales, molecular signatures suggesting the action of variable selective effects were observed along the genome. In contrast to the general tendency, some linkage groups displayed strong positive associations between the recombination rate and the domestic ancestry rate, which might be explained by several mechanisms such as: (i) local variation could reflect the stochastic outcomes of genetic drift (Martin & Jiggins, 2017). (ii) The presence of hybrid incompatibilities may also result in lower foreign ancestry in low recombining regions (Schumer et al., 2018). Such pattern has been observed in swordtail fish species where the retention of minor parent ancestry was more pronounced in highly recombining regions (Schumer et al., 2018), as well as in European sea bass (Duranton et al., 2018). Similarly, Martin et al. (2019) observed stronger barriers to introgression within longer chromosomes displaying lower average recombination rate on average. Finally, (iii) in the situation of selection against domestic haplotypes (i.e. deleterious introgression, or hybridization load), foreign loci of small individual effect are expected to be removed more quickly within low recombining regions because selection is more effective in removing linked deleterious mutations (Martin & Jiggins, 2017; Schumer et al., 2018; Kim et al., 2018). We observed such pattern for two LGs (LGs7 and 41) displaying a deficit of domestic introgression in the presence of potentially deleterious mutations. Moreover, LG7 broadly displayed lower domestic introgression within low recombining regions. Together, these combined pieces of information suggest that selection has been acting against the domestic haplotypes potentially carrying deleterious alleles.

### Importance of the time since hybridization

The time since hybridization seems to be an important parameter that determines the fate of admixture tracts, as different ancestry patterns were observed as a function of the hybrids category considered. Late-generation hybrids displayed a stronger negative correlation between domestic ancestry and recombination rate and even reversed pattern for three linkage groups (LGs 7, 19 and 34) compared to the early hybrid generations. These discrepancies between the early and late-generation hybrids could be caused by a time-specific effect of selection. Indeed, the outcomes of hybridization are expected to vary with the number of generations elapsed since the introduction of foreign alleles into the population. If the hybridization event is recent, long introgressed haplotypes are expected since it takes time for recombination to break down domestic tracts across generations (Racimo et al., 2015). Thus the selective effects in recent hybrids generations are expected to act at the scale of haplotype blocks, similarly as in low recombining regions. Inversely, smaller introgressed haplotypes are expected for older hybridization events (Racimo et al., 2015) and thus selective effects would act at a more localized (i.e. locus) scale, similarly as for high recombining regions (see above)(Leitwein et al., 2018; McFarlane & Pemberton, 2018). Here, we suspect that for the early hybrid individuals, the admixture events were too recent for selection to efficiently occur and be detectable. Continuously, in our late hybrids individuals, while the introgressed haplotypes are smaller than the early hybrids individuals they are still long (~15Mbp) and selection would tend to act at the block scale explaining the predominance of the associative overdominance effects. As a consequence, the hybridization events studied here are still too recent to fully assess de long term selective outcomes of supplementation. Indeed, while positive effect of associative overdominance are predominant in our study, it is possible that later on maladaptative introgressed domestic alleles reveal their individual effects, which could become detrimental for the supplemented populations (Harris & Nielsen, 2016).

### Implications for conservation biology

Understanding the evolutionary outcomes of anthropogenic hybridization is of major concerns for conservation biology and management (Allendorf, 2017; McFarlane & Pemberton, 2018). Indeed, supplementation with a foreign population could either be beneficial for the recipient population (i.e. genetic rescue; Allendorf et al., 2010; Uller & Leimu, 2011; Aitken & Whitlock, 2013; Goedbloed et al., 2013), or on contrary could induce outbreeding depression because of maladaptation, loss of local adaption or hybrids incompatibilities (Waples, 1991; Orr, 1995; Randi, 2008; Verhoeven et al., 2011). Here, quantifying the relationship between introgression and recombination rates allowed revealing complex patterns of selective effects acting along the genome. Such pattern revealed that interpreting the consequences of anthropogenic hybridization is not straightforward, since several antagonistic mechanisms may cause both positive and negative effects of domestic ancestry to co-occur along the genome. From a conservation point of view, it is important to weigh the pros and cons of those different outcomes and evaluate the conservation main focus (i.e. the species, the population or the local genetic inheritance). Our method may help to consider the potential evolutionary consequences of hybridization on the short and possibly long terms in order to take sound management decisions. Moreover, evaluating the length and distribution of introgressed haplotypes allows determining the approximate time since hybridization (Leitwein et al., 2018; Duranton, Bonhomme, & Gagnaire, 2019) and hybrid categories, which can influence management practices and conservation decisions (Allendorf, Leary, Spruell, & Wenburg, 2001). Indeed, a population carrying a small proportion of domestic ancestry might be of more interest from a conservation standpoint compared to a population carrying a stronger one. This seems particularly important when the history of supplementation is unknown, for instance due to illicit stocking (Johnson, Arlinghaus, & Martinez, 2009). Moreover, the selective forces modulating the introgression rate along the genome will be influenced by the length of introgressed haplotypes which is dependent on both the number of generations since hybridization and the recombination rate. As a result, both the time and the recombination rate variation are important to understand the consequences (positive or negative) of hybridization with a foreign population and better orient management decisions. Finally, applying such methods to non-model species are becoming more and more accessible and thus may be helpful in conservation for a wide range of natural hybridization contexts.

### Conclusions

It has recently been recognized that consideration of the recombination rate is of prime importance to interpret how natural selection shapes the genomic landscape of introgression in admixed populations (Martin & Jiggins, 2017; Duranton et al., 2018; Schumer et al., 2018; S. H. Martin et al., 2019). In our study we highlight the importance of the temporal dynamics of hybridization and the genome-wide variation of the recombination rates, both resulting in a complex interplay of multiple evolutionary processes occurring along the genome. By assessing the pattern of introgression and recombination at three different scales (i.e. global, linkage group and 2Mb windows size) we were able to provide a detailed picture of these antagonistic evolutionary mechanisms (e.g., positive and negative selection) occurring along the genome. In particular, our results show that the interplay between the recombination rate and the presence of potentially deleterious recessive mutations may be responsible for these variable selective patterns along the genome. Such variability of both selective forces and recombination rate along the genome reflect the complexity of genome evolution and need to be considered before drawing conclusions regarding the beneficial and/or negative effects of hybridization. Especially, since an apparent beneficial outcome of hybridization during the early generations could be detrimental in later hybrid generations when the individual effects of maladaptative loci are being exposed (Harris & Nielsen, 2016). In the same way, the whole genome tendency could display a general beneficial outcome of hybridization while analyses at the local scale could reveal strong selection against maladaptative introgressed alleles.

## Acknowledgments

We thank biologists and technicians of the Société d’Etablissement de Plain Air du Québec (SEPAQ) and the Ministère des Forêts, de la Faune et des Parcs du Québec (MFFP), in particular Amélie Gilbert and Isabel Thibault for their implication in the project and/or their field assistance and sampling. This research was funded by the MFFP, the Canadian Research Chair in Genomics and Conservation of Aquatic Resources, Ressources Aquatiques Québec (RAQ) as well as by a Strategic Project Grant from the Natural Science and Engineering Research Council of Canada (NSERC) to L. Bernatchez, D. Garant and P. Sirois.

## Data accessibility

Supporting Information Table Sup1 displays the individual chromosomal ancestry imbalance, the number of domestic haplotypes, the total percentage of domestic ancestry per individuals, the number of generations since the mean years of stocking practice and the hybrids generation. Table Sup2 describe the individual domestic ancestry tracts position and length. Raw data are available from Létourneau et al. (2018) at Dryad Digital Repository: https://doi.org/10.5061/dryad.s5qt3.

## Author contributions

M.L. and L.B. conceived the study. M.L. and H.C. designed the analyses and performed the analyses. E.N., H.C. and A-L.F. contribute to the bioinformatics and analyses interpretation. M.L. wrote the manuscript. P-A.G. helps with the manuscript structuration and all co-authors critically revised the manuscript and approved the final version to be published.

---

**Figure Sup. 1.**
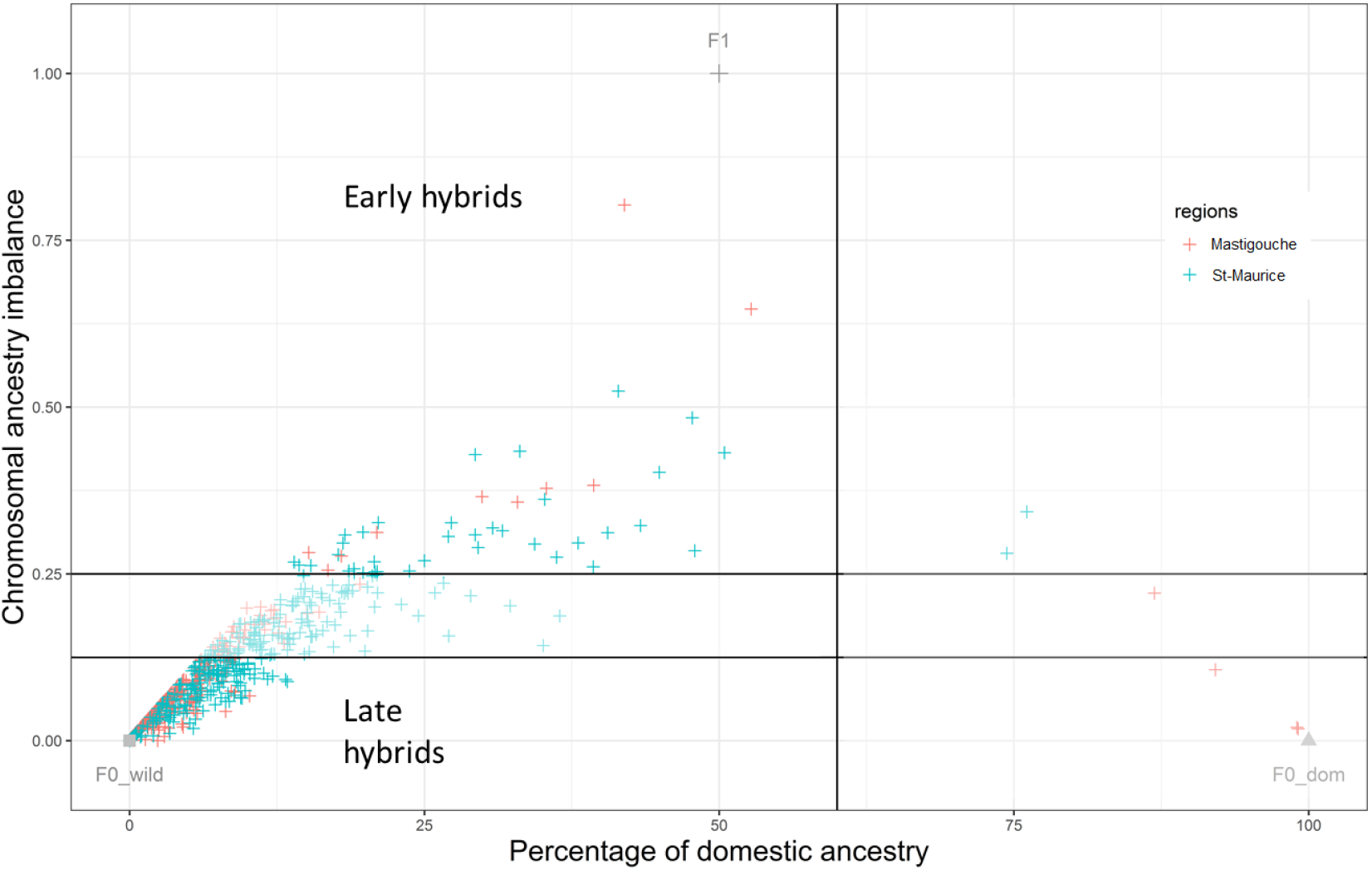
Plot of the chromosomal ancestry imbalance (CAI) as a function of the percentage of domestic ancestry for each wild-caught admixed individual considered. Grey points represent the theoretical expectations for F0_wild (without domestic ancestry, thus a CAI of 0 because individuals are theoretically pure wild), F0_dom (without wild ancestry, individuals are theoretically pure domestic (CAI= 0)) and F1 individuals (with half domestic ancestry and half wild ancestry, thus the CAI is maximum (CAI = 1) between homologues). The colours represent the two reserves: red for Mastigouche and green for the St Maurice. Horizontal and perpendicular lines represent the delineation of both early and late hybrid generations.

**Figure Sup. 2.**
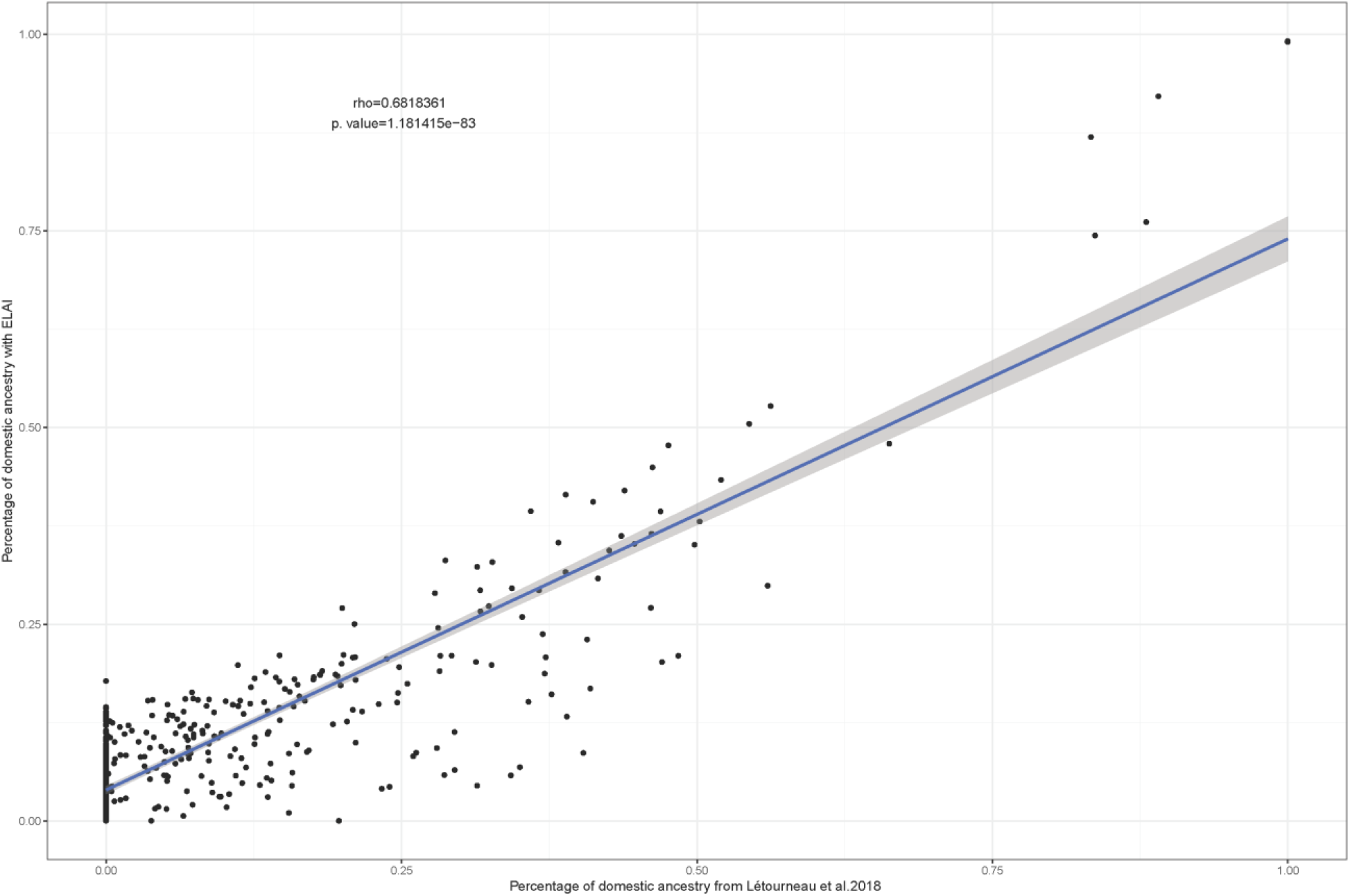
Positive correlation between the mean individual domestic ancestries estimated with with ELAI and with ADMIXTURE from Létoureau et al. (2018) (rho spearman’s = 0.68, p-value<0.001).

**Figure Sup. 3.**
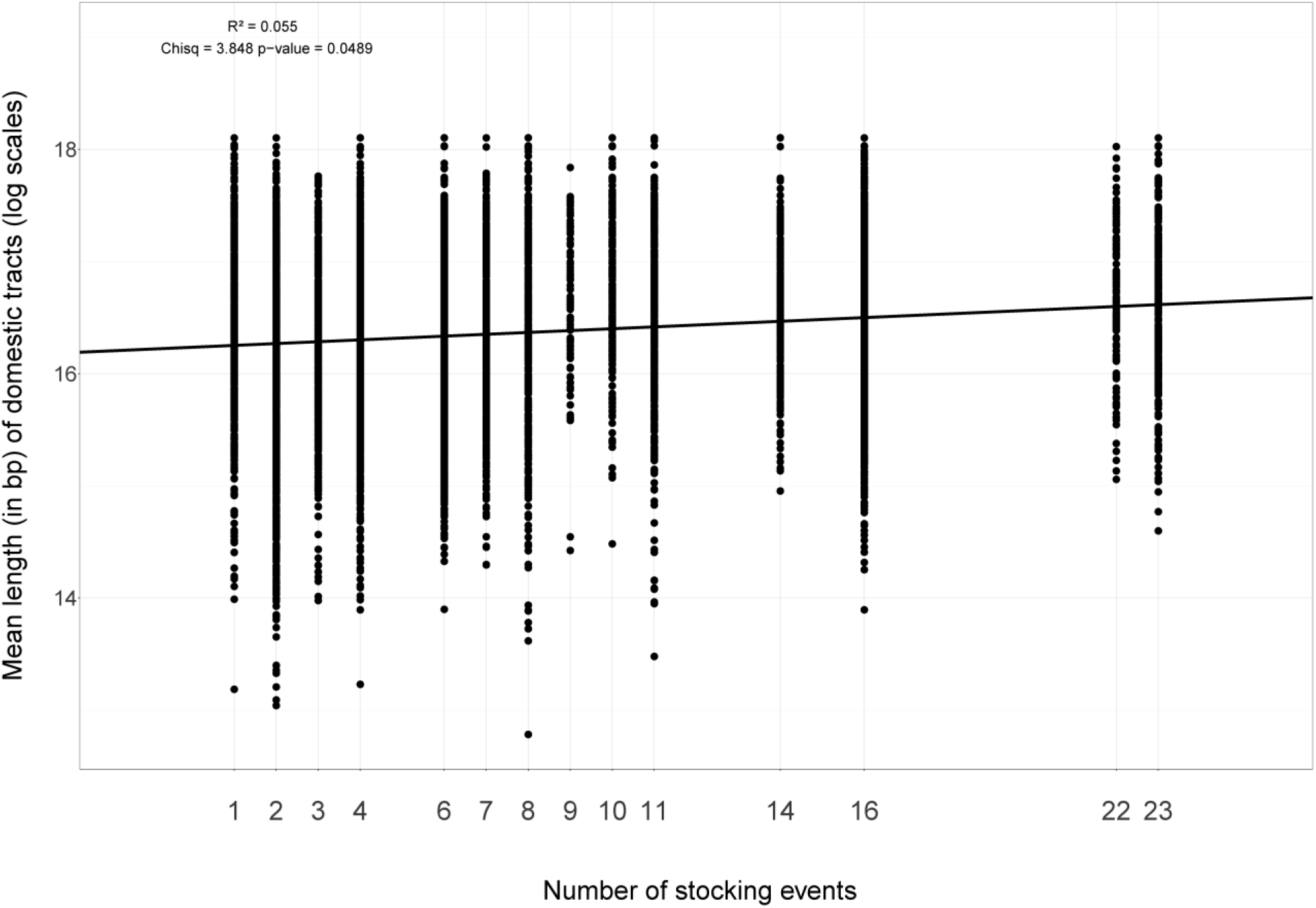
Positive correlation between the mean length (in bp log scales) of domestic tracts per individual as a function of the number of stocking events for each sampled lake (lakes labels are described in table 1; R^2^=0.055, p-value=0.0489).

**Figure Sup. 4.**
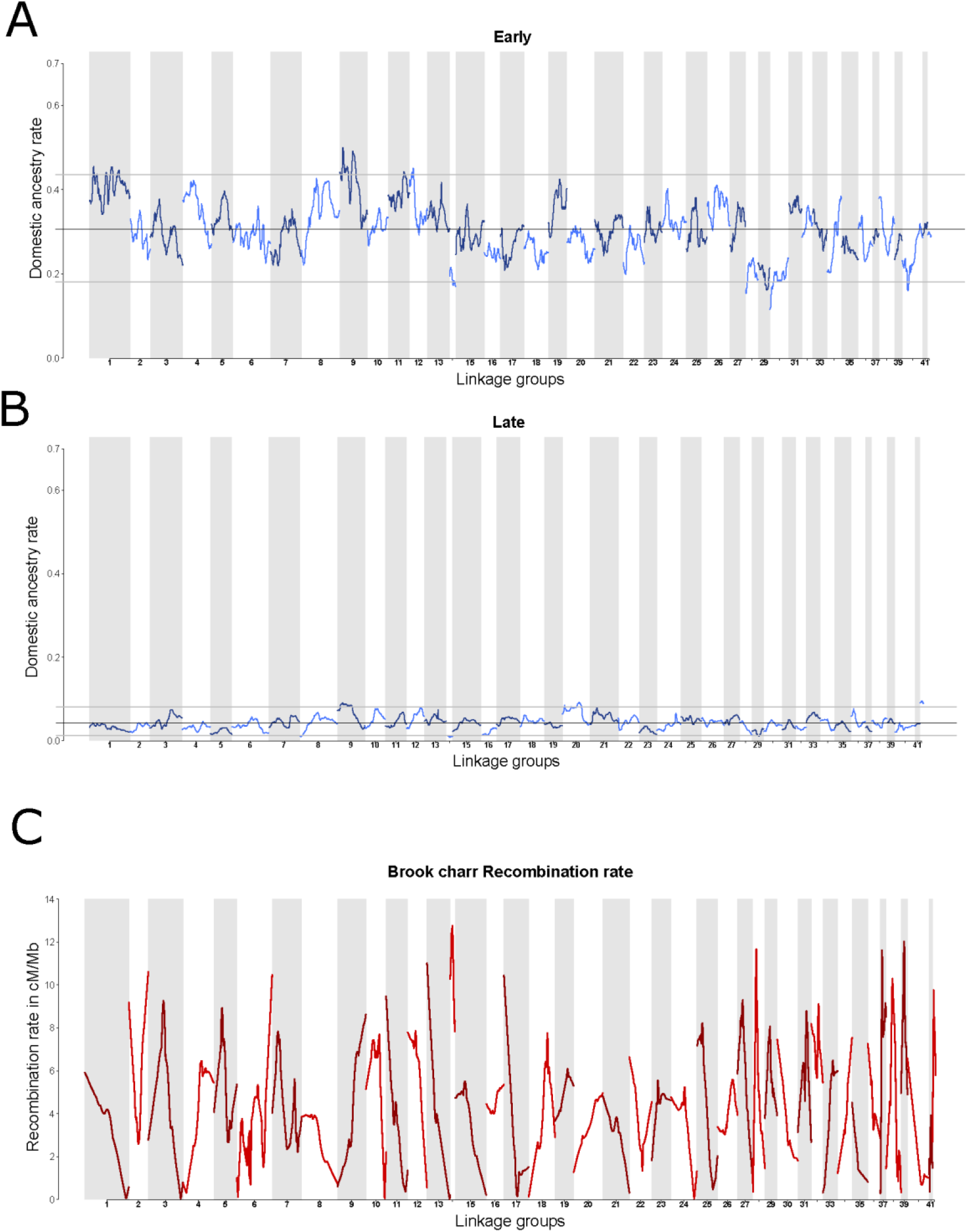
Genome-wide introgression rate of the domestic ancestry for the early (A) and late (B) hybrids along the 42 Brook Charr linkage groups (LGs). Black line represents the mean introgression rate and gray lines the 95% confidence intervals. C) Estimates of the Brook Charr recombination rates in cM/Mb along the 42 LGs.

**Figure Sup. 5.**
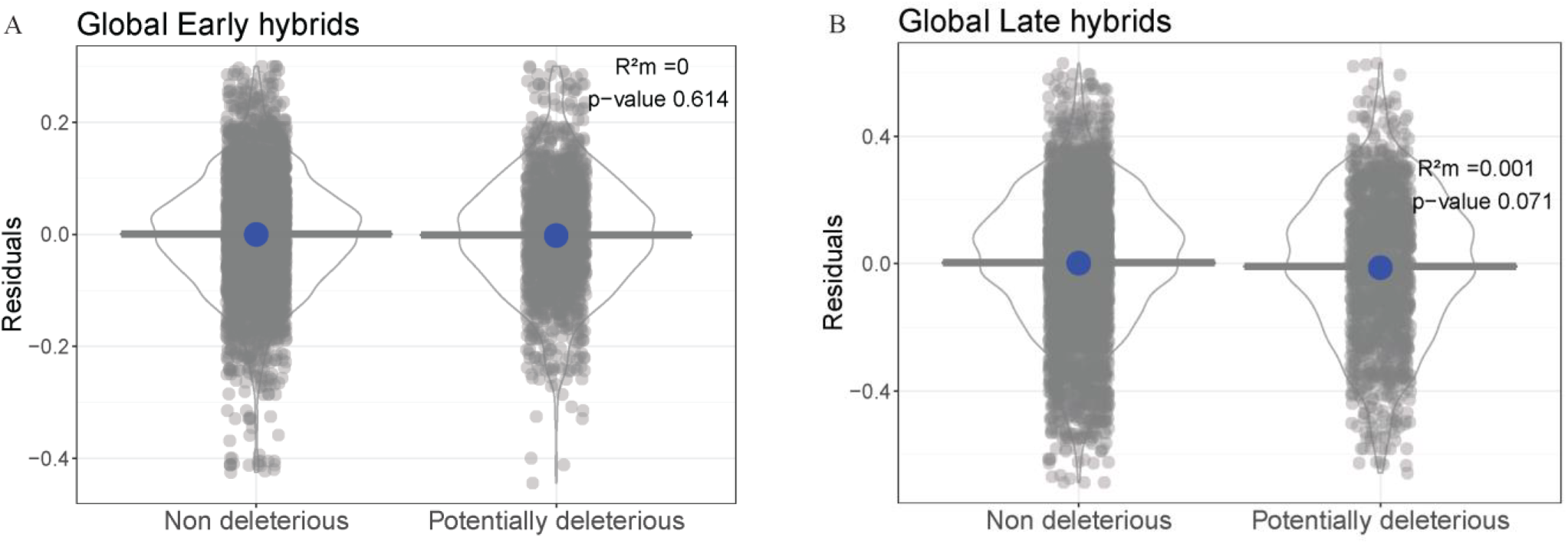
Residuals of the regression between the domestic ancestry and recombination rates models as a function of the presence of non-deleterious or potentially deleterious alleles at the genome-wide scale for the Early (A) and Late-generation hybrids(B). Horizontal line represents the median. Blue dots and blue line represent the estimates and the confidence intervals of the linear model estimates. Marginal R^2^ (R^2^m) and significance are displayed at the upper right of each plot.

## Reference

Aitken, S. N., & Whitlock, M. C. (2013). Assisted Gene Flow to Facilitate Local Adaptation to Climate Change. Annual Review of Ecology, Evolution, and Systematics, 44(1), 367–388. doi: 10.1146/annurev-ecolsys-110512-135747

Allendorf, F. W. (2017). Genetics and the conservation of natural populations: allozymes to genomes. Molecular Ecology, 26(2), 420–430. doi: 10.1111/mec.13948

Allendorf, F. W., Hohenlohe, P. A., & Luikart, G. (2010). Genomics and the future of conservation genetics. Nature Reviews Genetics, 11(10), 697–709. doi: 10.1038/nrg2844

Allendorf, F. W., Leary, R. F., Spruell, P., & Wenburg, J. K. (2001). The problems with hybrids: setting conservation guidelines. Trends in Ecology & Evolution, 16(11), 613–622. doi: 10.1016/S0169-5347(01)02290-X

Anderson, E., & Stebbins, G. L. (1954). Hybridization as an Evolutionary Stimulus. Evolution, 8(4), 378–388. doi: 10.1111/j.1558-5646.1954.tb01504.x

Bates, D., Mächler, M., Bolker, B., & Walker, S. (2015). Fitting Linear Mixed-Effects Models Using lme4. Journal of Statistical Software, 67(1), 1–48. doi: 10.18637/jss.v067.i01

Catchen, J., Hohenlohe, P. A., Bassham, S., Amores, A., & Cresko, W. A. (2013). Stacks: an analysis tool set for population genomics. Molecular Ecology, 22(11), 3124–3140.

Charlesworth, B. (2009). Effective population size and patterns of molecular evolution and variation. Nature Reviews Genetics, 10(3), 195–205. doi: 10.1038/nrg2526

Charlesworth, D., & Willis, J. H. (2009). The genetics of inbreeding depression. Nature Reviews Genetics, 10(11), 783–796. doi: 10.1038/nrg2664

Chen, Z. J. (2010). Molecular mechanisms of polyploidy and hybrid vigor. Trends in Plant Science, 15(2), 57–71. doi: 10.1016/j.tplants.2009.12.003

Choi, Y., Sims, G. E., Murphy, S., Miller, J. R., & Chan, A. P. (2012). Predicting the Functional Effect of Amino Acid Substitutions and Indels. PLOS ONE, 7(10), e46688. doi: 10.1371/journal.pone.0046688

Christensen, K. A., Rondeau, E. B., Minkley, D. R., Leong, J. S., Nugent, C. M., Danzmann, R. G., … Koop, B. F. (2018). The Arctic charr (Salvelinus alpinus) genome and transcriptome assembly. PLOS ONE, 13(9), e0204076. doi: 10.1371/journal.pone.0204076

Dannemann, M., & Racimo, F. (2018). Something old, something borrowed: Admixture and adaptation in human evolution (No. e26782v2). doi: 10.7287/peerj.preprints.26782v2

Duranton, M., Allal, F., Fraïsse, C., Bierne, N., Bonhomme, F., & Gagnaire, P.-A. (2018). The origin and remolding of genomic islands of differentiation in the European sea bass. Nature Communications, 9(1), 2518. doi: 10.1038/s41467-018-04963-6

Duranton, M., Bonhomme, F., & Gagnaire, P.-A. (2019). The spatial scale of dispersal revealed by admixture tracts. Evolutionary Applications, 0(ja). doi: 10.1111/eva.12829

Fay, J. C., & Wu, C.-I. (2000). Hitchhiking Under Positive Darwinian Selection. Genetics, 155(3), 1405–1413.

Felsenstein, J. (1974). The Evolutionary Advantage of Recombination. Genetics, 78(2), 737–756.

Ferchaud, A.-L., Laporte, M., Perrier, C., & Bernatchez, L. (2018). Impact of supplementation on deleterious mutation distribution in an exploited salmonid. Evolutionary Applications, 11(7), 1053–1065. doi: 10.1111/eva.12660

Ferchaud, A.-L., Leitwein, M., Laporte, M., Boivin-Delisle, D., Bougas, B., Hernandez, C., … Bernatchez, L. (2019). Adaptive and maladaptive genetic diversity in small populations; insights from the Brook Charr (Salvelinus fontinalis) case study | bioRxiv. Retrieved from https://www.biorxiv.org/content/10.1101/660621v1.abstract

Frankham, R. (2015). Genetic rescue of small inbred populations: meta-analysis reveals large and consistent benefits of gene flow. Molecular Ecology, 24(11), 2610–2618. doi: 10.1111/mec.13139

Goedbloed, D. J., van Hooft, P., Megens, H.-J., Langenbeck, K., Lutz, W., Crooijmans, R. P., … Prins, H. H. (2013). Reintroductions and genetic introgression from domestic pigs have shaped the genetic population structure of Northwest European wild boar. BMC Genetics, 14, 43. doi: 10.1186/1471-2156-14-43

Gosselin, T., & Bernatchez, L. (2016). Stackr: GBS/RAD data exploration, manipulation and visualization using R. Retrieved from https://github.com/thierrygosselin/stackr. Retrieved from https://github.com/thierrygosselin/stackr

Gossieux, P., Bernatchez, L., Sirois, P., & Garant, D. (2019). Impacts of stocking and its intensity on effective population size in Brook Charr (Salvelinus fontinalis) populations. Conservation Genetics.

Gravel, S. (2012). Population Genetics Models of Local Ancestry. Genetics, 191(2), 607–619. doi: 10.1534/genetics.112.139808

Guan, Y. (2014). Detecting Structure of Haplotypes and Local Ancestry. Genetics, 196(3), 625–642. doi: 10.1534/genetics.113.160697

Harris, K., & Nielsen, R. (2013). Inferring Demographic History from a Spectrum of Shared Haplotype Lengths. PLOS Genet, 9(6), e1003521. doi: 10.1371/journal.pgen.1003521

Harris, K., & Nielsen, R. (2016). The Genetic Cost of Neanderthal Introgression. Genetics, 203(2), 881–891. doi: 10.1534/genetics.116.186890

Harris, K., Zhang, Y., & Nielsen, R. (2019). Genetic rescue and the maintenance of native ancestry. Conservation Genetics. doi: 10.1007/s10592-018-1132-1

Huerta-Sánchez, E., Jin, X., Asan, Bianba, Z., Peter, B. M., Vinckenbosch, N., … Nielsen, R. (2014). Altitude adaptation in Tibetans caused by introgression of Denisovan-like DNA. Nature, 512(7513), 194–197. doi: 10.1038/nature13408

Johnson, B. M., Arlinghaus, R., & Martinez, P. J. (2009). Are We Doing All We Can to Stem the Tide of Illegal Fish Stocking? Fisheries, 34(8), 389–394. doi: 10.1577/1548-8446-34.8.389

Kim, B. Y., Huber, C. D., & Lohmueller, K. E. (2018). Deleterious variation shapes the genomic landscape of introgression. PLOS Genetics, 14(10), e1007741. doi: 10.1371/journal.pgen.1007741

Lamaze, F. C., Sauvage, C., Marie, A., Garant, D., & Bernatchez, L. (2012). Dynamics of introgressive hybridization assessed by SNP population genomics of coding genes in stocked brook charr (Salvelinus fontinalis). Molecular Ecology, 21(12), 2877–2895.

Leitwein, M., Gagnaire, P.-A., Desmarais, E., Berrebi, P., & Guinand, B. (2018). Genomic consequences of a recent three-way admixture in supplemented wild brown trout populations revealed by local ancestry tracts. Molecular Ecology, 27(17), 3466–3483. doi: 10.1111/mec.14816

Létourneau, J., Ferchaud, A.-L., Luyer, J. L., Laporte, M., Garant, D., & Bernatchez, L. (2018). Predicting the genetic impact of stocking in Brook Charr (Salvelinus fontinalis) by combining RAD sequencing and modeling of explanatory variables. Evolutionary Applications, 11(5), 577–592. doi: 10.1111/eva.12566

Li, H., & Durbin, R. (2010). Fast and accurate long-read alignment with Burrows–Wheeler transform. Bioinformatics, 26(5), 589–595. doi: 10.1093/bioinformatics/btp698

Lippman, Z. B., & Zamir, D. (2007). Heterosis: revisiting the magic. Trends in Genetics, 23(2), 60–66. doi: 10.1016/j.tig.2006.12.006

Martin, S. H., Davey, J., Salazar, C., & Jiggins, C. D. (2019). Recombination rate variation shapes barriers to introgression across butterfly genomes. doi: https://doi.org/10.1371/journal.pbio.2006288

Martin, S., & Jiggins, C. D. (2017). Interpreting the genomic landscape of introgression. Current Opinion in Genetics & Development, 47, 69–74. doi: 10.1016/j.gde.2017.08.007

Martin, S., Savaria, J.-Y., Audet, C., & Bernatchez, L. (1997). Microsatellites reveal no evidence for inbreeding effects but low inter-stock genetic diversity among brook charr stocks used for production in Québec. 97(2)(21–23).

McFarlane, E., & Pemberton, J. M. (2018). Detecting the True Extent of Introgression during Anthropogenic Hybridization: Trends in Ecology & Evolution. Retrieved from https://www.cell.com/trends/ecology-evolution/fulltext/S0169-5347(18)30305-7?_returnURL=https%3A%2F%2Flinkinghub.elsevier.com%2Fretrieve%2Fpii%2FS0169534718303057%3Fshowall%3Dtrue

Nugent, C. M., Easton, A. A., Norman, J. D., Ferguson, M. M., & Danzmann, R. G. (2017). A SNP Based Linkage Map of the Arctic Charr (Salvelinus alpinus) Genome Provides Insights into the Diploidization Process After Whole Genome Duplication. G3: Genes, Genomes, Genetics, 7(2), 543–556. doi: 10.1534/g3.116.038026

Orr, H. A. (1995). The population genetics of speciation: the evolution of hybrid incompatibilities. Genetics, 139(4), 1805–1813.

Racimo, F., Sankararaman, S., Nielsen, R., & Huerta-Sánchez, E. (2015). Evidence for archaic adaptive introgression in humans. Nature Reviews Genetics, 16(6), 359–371. doi: 10.1038/nrg3936

Randi, E. (2008). Detecting hybridization between wild species and their domesticated relatives. Molecular Ecology, 17(1), 285–293. doi: 10.1111/j.1365-294X.2007.03417.x

Renaut, S., & Rieseberg, L. H. (2015). The Accumulation of Deleterious Mutations as a Consequence of Domestication and Improvement in Sunflowers and Other Compositae Crops. Molecular Biology and Evolution, 32(9), 2273–2283. doi: 10.1093/molbev/msv106

Rezvoy, C., Charif, D., Guéguen, L., & Marais, G. A. (2007). MareyMap: an R-based tool with graphical interface for estimating recombination rates. Bioinformatics, 23(16), 2188–2189.

Schumer, M., Xu, C., Powell, D. L., Durvasula, A., Skov, L., Holland, C., … Przeworski, M. (2018). Natural selection interacts with recombination to shape the evolution of hybrid genomes. Science, 360(6389), 656–660. doi: 10.1126/science.aar3684

Sutherland, B. J. G., Gosselin, T., Normandeau, E., Lamothe, M., Isabel, N., Audet, C., & Bernatchez, L. (2016). Salmonid chromosome evolution as revealed by a novel method for comparing RADseq linkage maps. Genome Biology and Evolution, evw262. doi: 10.1093/gbe/evw262

Team, R. C. (2015). R: A language and environment for statistical computing [Internet]. Vienna, Austria: R Foundation for Statistical Computing; 2013. Document Freely Available on the Internet at: Http://Www.r-Project.Org.

Todesco, M., Pascual, M. A., Owens, G. L., Ostevik, K. L., Moyers, B. T., Hübner, S., … Rieseberg, L. H. (2016). Hybridization and extinction. Evolutionary Applications, 9(7), 892–908. doi: 10.1111/eva.12367

Turelli, M., & Orr, H. A. (2000). Dominance, Epistasis and the Genetics of Postzygotic Isolation. Genetics, 154(4), 1663–1679.

Uller, T., & Leimu, R. (2011). Founder events predict changes in genetic diversity during human-mediated range expansions. Global Change Biology, 17(11), 3478–3485. doi: 10.1111/j.1365-2486.2011.02509.x

Verhoeven, K. J. F., Macel, M., Wolfe, L. M., & Biere, A. (2011). Population admixture, biological invasions and the balance between local adaptation and inbreeding depression. Proceedings of the Royal Society of London B: Biological Sciences, 278(1702), 2–8. doi: 10.1098/rspb.2010.1272

Waples, R. S. (1991). Genetic interactions between hatchery and wild salmonids: lessons from the Pacific Northwest. Canadian Journal of Fisheries and Aquatic Sciences, 48(S1), 124–133.

